# Ovarian development is driven by early spatiotemporal priming of the coelomic epithelium

**DOI:** 10.1101/2025.09.24.678234

**Authors:** Cyril Djari, Chloé Mayère, Maëva Guy, Aitana Perea-Gomez, Laura Bellutti, Paul Barreau, Herta Ademi, Agathe Rozier, Anthony S. Martinez, Tyler J. Gibson, Françoise Kühne, Cassandre Guérin, Dagmar Wilhelm, Gabriel Livera, Jennifer McKey, Marie-Christine Chaboissier, Serge Nef

## Abstract

Ovarian organogenesis relies on the coordinated specification of supporting and steroidogenic lineages from multipotent coelomic epithelium (CE) progenitors. However, it remains unclear how ovarian cellular diversity arises and whether CE progenitors are fate-biased before or after ingression into the gonad. We show that CE cells in fetal mouse and human ovaries are transcriptionally heterogeneous and spatially organized into subdomains already primed toward supporting or steroidogenic identities. CE priming is dynamic, influenced by proximity to the mesonephros, with a transient coexistence of both progenitor types before resolution toward a predominantly supporting-biased CE. In mice, delamination of primed CE cells seeds intragonadal niches that generate pre-granulosa and steroidogenic progenitors. We further demonstrate that fetal steroidogenic progenitors give rise to adult stromal and theca cells, and that granulosa cells have a dual origin from CE-derived and supporting-like cells. Together, these findings reveal a conserved, spatially encoded program of ovarian lineage specification.

## Introduction

The sex of an individual is established at fertilization by the paternal contribution of either an X or a Y chromosome, resulting in an XX or XY embryo (Saunders and Veyrunes, 2021). This genetic sex directs the differentiation of the initially bipotential gonads into ovaries in XX embryos or testes in XY embryos. This process, known as gonadal sex determination, is fundamental to sexual development and fertility, as hormones secreted by the differentiated gonads subsequently regulate the development of primary and secondary sexual characteristics as well as gametogenesis. Sex determination occurs at 5–7 weeks in humans and at embryonic day (E)11.5 in mice, with a slight delay in XX embryos compared with XY embryos (Stevant et al., 2019; Wilhelm et al., 2025). Prior to sex determination, proliferation of multipotent coelomic epithelium (CE) progenitors in the emerging gonadal primordium is essential for gonadal formation (Karl and Capel, 1998; Mayere et al., 2022; Schmahl et al., 2000). CE progenitors divide asymmetrically, with some daughter cells remaining epithelial while others undergo epithelial-to-mesenchymal transition (EMT) and ingress into the underlying gonadal tissue. This process drives gonadal growth and establishes the somatic foundation of the developing gonad (Karl and Capel, 1998; Kusaka et al., 2010; Lin et al., 2017; Schmahl et al., 2000).

Multipotent CE progenitors generate most somatic cells in both XX and XY gonads and give rise to the two major somatic lineages of the gonad: the supporting and steroidogenic lineages (Ademi et al., 2022; DeFalco et al., 2011; Karl and Capel, 1998; Liu et al., 2015; Liu et al., 2016; Mayere et al., 2022; Mork et al., 2012; Schmahl et al., 2003; Stevant and Nef, 2018, 2019). In XX gonads, these lineages form pre-granulosa and theca/stromal cells, whereas in XY gonads, they generate Sertoli and Leydig cells, respectively. Despite their central importance, the mechanisms underlying CE progenitor diversification remain incompletely understood. A key unresolved question is whether the CE contains heterogeneous progenitor populations already biased toward supporting or steroidogenic fates, or whether lineage specification occurs only after delamination into the gonad. If such heterogeneity exists, how these progenitors are temporally regulated and spatially organized to generate the two somatic lineages remains unknown.

Another major unresolved question in ovarian biology is how granulosa and steroidogenic lineages arise during development. Pre-granulosa cells emerge in two waves, a medullary first wave (Mork et al., 2012; Niu and Spradling, 2020; Zheng et al., 2014) and a cortical second wave (Cai et al., 2020; Feng et al., 2016; Niu and Spradling, 2020; Rastetter et al., 2014; Stevant et al., 2019), but the dynamics and relative contributions of these populations remain unclear, particularly given recent evidence that XX supporting-like cells (SLCs) also generate a subset of pre-granulosa cells (Mayere et al., 2022; McKey et al., 2022). These pre-granulosa cells subsequently differentiate into granulosa cells at the onset of folliculogenesis after birth in mice (Mork et al., 2012; Zheng et al., 2014). Similarly, although theca cells differentiate only after birth (Liu et al., 2015; Magoffin and Weitsman, 1994), the specification and maintenance of their fetal progenitors remain poorly defined. We refer to these cells as stromal progenitors (SPs), a population encompassing both steroidogenic progenitors and non-steroidogenic stromal derivatives. How these distinct progenitor pools are specified, spatially organized, and integrated into ovarian development therefore remains a central open question.

To address how ovarian cellular diversity arises upstream of sex determination and how granulosa and steroidogenic lineages first emerge, we combined single-cell transcriptomics in mouse and human with spatial transcriptomics, *in vivo* lineage tracing, and *ex vivo* functional assays. These approaches reveal that granulosa cells have a dual origin and that spatially patterned CE subpopulations are primed toward distinct fates and directly seed ovarian lineages through localized delamination, establishing intragonadal niches with important implications for ovarian development.

## Results

### Coelomic epithelium and supporting-like cells are conserved progenitors of ovarian somatic lineages in humans and mice

To investigate the specification and differentiation dynamics of supporting and stromal/steroidogenic lineages in humans and mice, we generated species-specific single-cell RNA-sequencing (scRNA-seq) atlases of ovarian development spanning post-conception weeks (PCW) 5-21 in humans and embryonic days (E)10.5 to post-natal day (P)56 in mice. These atlases were assembled from two human-focused studies (Garcia-Alonso et al., 2022; Lardenois et al., 2026) and three mouse single-cell datasets (Mayère et al., 2022; Niu and Spradling, 2020; Rossitto et al., 2022) (**Fig. S1A-F**). After annotation harmonization and selection of ovary somatic cells based on *GATA4*/*Gata4* expression, we generated two refined somatic sub-atlases comprising 156,408 human cells **(Fig 1A, B** and **Fig. S1C**) and 43,053 mouse cells **(Fig 1C, D** and **Fig. S1D**). In both species, these sub-atlases included coelomic and surface epithelial cells (CE/SE; *UPK3B* /*Upk3b*^+^), supporting-like cells (SLCs; *PAX8* /*Pax8*^+^), stromal progenitors (SPs; *PDGFRA* in human/*Nr2f2*^+^ in mouse), as well as pre-granulosa cells. In humans, pre-granulosa cells segregated into two distinct sub-populations corresponding to the first- and second-wave pre-granulosa cells (preGran-I; *CYP19A1*^+^ and preGran-II; *IRX3*^+^, *LHX2^+^*) **(Fig. S1E**). Mouse pre-granulosa cells formed a continuous cell community merging first-wave (*Foxl2^+^*) and second-wave (*Lgr5^+^*) pre-granulosa (**Fig. S1F**).

**Figure 1.**
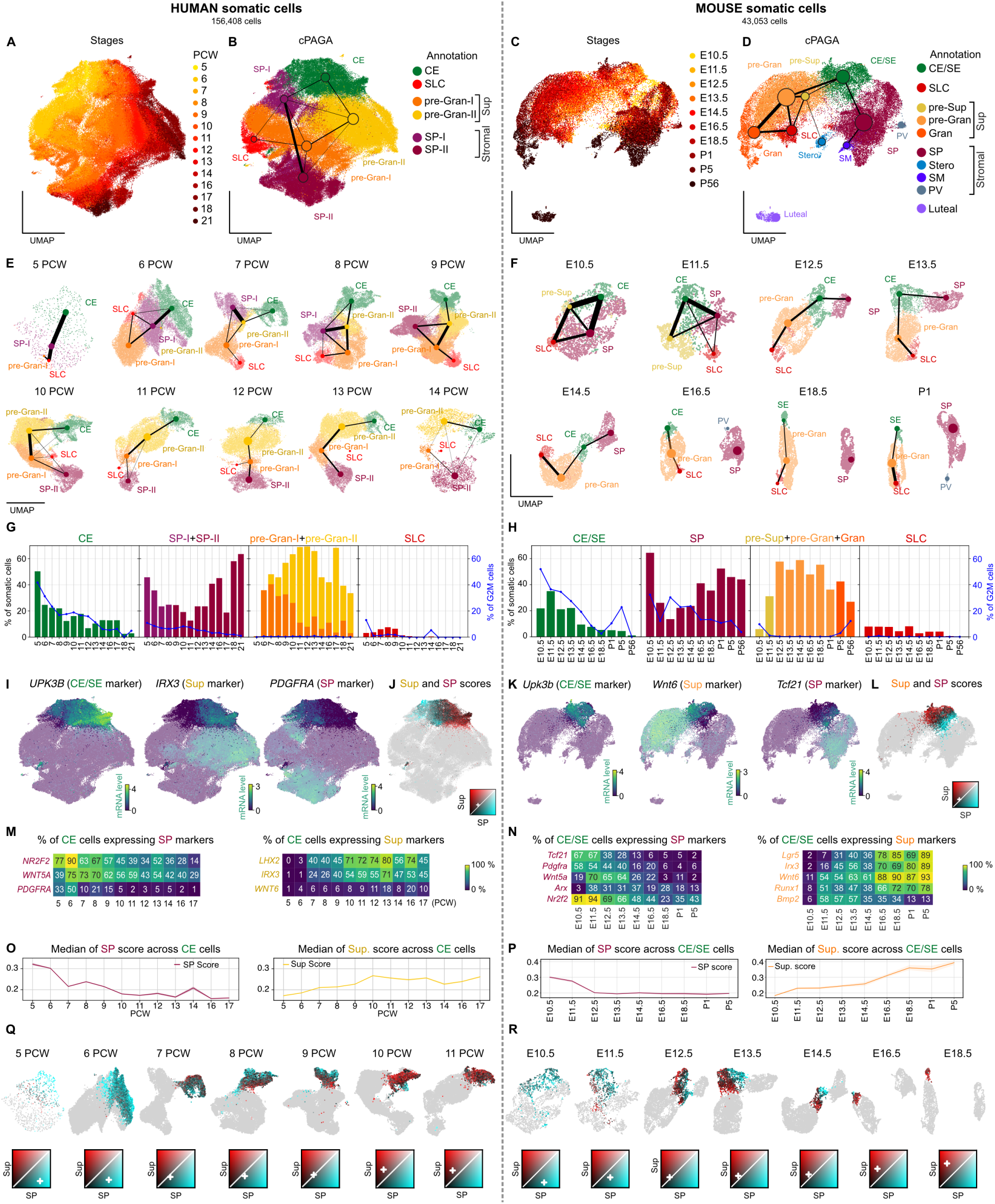
Single-cell transcriptomic atlas of the developing human and mouse ovaries reveals fate-biased heterogeneity in the coelomic epithelium and dual origin of granulosa cells. (A-D) UMAP representations of single somatic cells from human (n = 156,408) and mouse (n = 43,053) datasets. Human cells are colored by post-conceptional week (PCW) (A) and cell annotation (B); mouse cells are colored by embryonic and postnatal day (C) and cell annotation (D). In the annotated UMAPs, Conductance-based Partition-based Graph Abstraction (cPAGA) is overlaid to depict connectivity between somatic cell populations and across developmental stages. Edge width reflects the conductance score. Cell type abbreviations: CE, coelomic epithelium; SE, surface epithelium; SLC, supporting-like cells; pre-Sup, pre-supporting cells; pre-Gran, pre-granulosa cells; Gran, granulosa cells; SP, stromal progenitors; Stero, steroidogenic cells; SM, smooth muscle cells; PV, perivascular cells; Luteal, luteal cells. (E and F) Stage-wise UMAP representations colored by cell type, with cPAGA connectivity between cell types in human (E) and mouse (F). (G-H) Bar plots (left Y-axis) showing the proportion of CE/SE, pre-Gran, SP, and SLC populations across developmental stages in human (G) and mouse (H). Blue line plots (right Y-axis) indicate the proportion of proliferating cells (G2/M phase) within each population. (I and K) UMAPs of human (I) and mouse (K) somatic cells colored by expression of CE markers (*UPK3B/Upk3b*), supporting markers (*IRX3/Wnt6*), and SP markers (*PDGFRA/Tcf21*). (J and L) UMAP representations of human (J) and mouse (L) somatic cells. CE/SE cells are colored by scaled SP (turquoise channel) and supporting (red channel) identity scores. Legend squares indicate color channels, with SP scores mapped to the X-axis and supporting scores to the Y-axis. White diagonal lines indicate equivalent SP and supporting scores. White crosses represent the median score within CE cells. (M and N) Heatmap showing the proportion of cells expressing SP or supporting markers genes in the CE/SE population across developmental stages in human (M) and mouse (N). (O and P) Lineplot showing median supporting and SP identity scores with 95% confidence intervals across developmental stages in human (O) and mouse (P). (Q and R) Stage-specific UMAP representations of human (Q) and mouse (R) somatic cells. CE/SE cells are colored according to SP (turquoise channel) and supporting (red channel) identity scores. Legend squares indicate color channels as above. White diagonal lines represent equivalent identity scores; white crosses mark medians.

Partition-Based Graph Abstraction with conductance scoring (cPAGA) revealed that, in both species, CE progenitors form a central developmental node strongly connected to SPs and second-wave pre-granulosa cells (preGran-II in humans; *Lgr5* pre-granulosa cells in mice) (**Fig. 1B, D** and **FigS1F**). In contrast, SLCs showed a preferential connectivity with first-wave pre-granulosa cells (pre-Gran-I in humans and *Foxl2*^+^ pre-granulosa cells in mice). Stage-wise UMAP projection and cPAGA analyses further revealed a temporally ordered progression of the lineage relationships. In humans, CE cells were initially transcriptionally connected to SPs at 5-6 PCW, then transiently connected to both SPs and preGran-II during a dual phase at 7-9 PCW, before becoming exclusively connected to the supporting lineage from 10 PCW onward (**Fig. 1E** and **Fig. S1G**). In mouse, CE/SE cells were connected to both SPs and pre-supporting/supporting cell at early stages (E10.5 to E14.5), followed by exclusive CE–supporting connectivity from E16.5 onward, consistent with a dynamic progenitor role of CE/SE cells (**Fig. 1F** and **Fig. S1H**).

Cell proportion and proliferation analyses further supported this model (**Fig. 1G** and **H**). Despite high proliferative activity at early stages, CE/SE cells progressively declined in abundance over development. In contrast, the largely non-proliferative pre-Granulosa population expanded rapidly in both species, consistent with expansion driven predominantly by continuous differentiation and recruitment of CE-derived progenitors rather than by intrinsic proliferation. SPs exhibited complementary dynamics: following a possible initial contribution from CE/SE, they showed intermediate proliferation activity and an autonomous expansion, with proportions decreasing at early stages (PCW5-11 and E10.5-E12.5), before increasing again at later time points. SLCs display minimal proliferative activity and constitute a small population whose abundance progressively decreases over developmental time.

Altogether, these comparative analyses identify proliferative CE cells as a central progenitor pool for both supporting and stromal/steroidogenic lineages, contributing substantially to the expanding but largely non-proliferative pre-granulosa population, while SLCs represent a parallel progenitor source for pre-granulosa cells.

### Coelomic epithelium progenitors are transcriptionally heterogeneous and dynamically biased toward supporting or stromal fates

To determine whether CE progenitor cells (*UPK3B^+^/Upk3b^+^*) are initially homogeneous or instead contain fate-biased subpopulations, we examined the expression of established markers of pre-granulosa cells (*LHX2, BMP2/Bmp2, IRX3/Irx3, Lgr5, Runx1* and *Wnt6*) and stromal progenitors (*PDGFRA/Pdgfra*, *NR2F2/Nr2f2, WNT5A/Wnt5a, Arx* and *Tcf21*) (**Fig. 1I, K**, **Fig****.S1E** and **F**). In both species, CE cells display heterogeneous expression of supporting and stromal markers, revealing that the CE is not a uniform pool of progenitors. To assess whether this observation reflects the expression of a limited set of markers or instead indicates a broader transcriptional bias of CE progenitors toward a specific fate, we calculated supporting and stromal identity scores for each CE/SE cell. These scores were generated using regularized regression to identify the optimal set of transcripts that distinguishes supporting and SP populations from others, using differentiated supporting and SPs from PCW8 to 21 in humans and E12.5 to P5 in mice as reference populations (**Fig. S1I** and **J**). UMAP projection colored with supporting and SP identity scores revealed distinct CE/SE transcriptional states in both species: supporting-biased CE/SE cells with high supporting scores localized near the supporting cells, whereas SP-biased CE/SE cells with high SP scores clustered near the SP population (**Fig. 1J** and **L**).

We next examined whether CE transcriptional heterogeneity follows similar temporal dynamics in humans and mice. Initially, a high proportion of CE/SE cells expressed stromal-associated markers (*NR2F2/Nr2f2*, *WNT5A/Wnt5a*, *PDGFRA/Pdgfra*, *Arx* and *Tcf21)*, which progressively declined, while the proportion of cells expressing supporting associated markers (*LHX2, IRX3/Irx3, WNT6/Wnt6, Bmp2, Lgr5* and *Runx1*) followed an opposite pattern (**Fig. 1M** and **N**). Identity score analysis confirmed parallel developmental trajectories. Median SP identity scores peaked around the time of sex determination and declined rapidly thereafter (PCW5–10 in humans; E10.5–E12.5 in mice), whereas supporting scores increased from the beginning of sex determination (PCW6–10 in humans; E11.5–P5 in mice) (**Fig 1O** and **P**). Stage-specific UMAPs visualization further revealed that these fate biases are largely mutually exclusive at the single-cell level. In humans, CE cells progressed from a predominantly SP-biased state at PCW5-6, then entered a transient dual phase at PCW7–9, characterized by distinct subpopulations biased toward either supporting or SP fates, after which CE cells became predominantly supporting-biased from PCW10 onward (**Fig 1Q**). Mouse CE/SE dynamics closely mirrored this three-phase progression: CE cells initially displayed a stronger SP bias, then distinct supporting- and SP-biased groups at E12.5-E14.5 and finally became predominantly supporting-biased from E16.5 onward (**Fig 1R**).

Together, these analyses reveal that CE is transcriptionally heterogeneous and follows a conserved, temporally ordered program. In both humans and mice, CE progenitors transition from an early phase dominated by SP-biased cells, to a transitional phase characterized by mutually exclusive subpopulations biased toward either supporting or SP fates, to a final supporting-restricted phase, progressively refining lineage potential while preserving supporting fate competence.

### Spatial patterning of the ovarian epithelium defines biased surface domains

Having established the conserved temporal dynamics of CE transcriptomic biases in humans and mice, we next asked whether these biased states are spatially organized along the ovarian surface during development. We focused our spatial analyses on the mouse ovary, whose small size and ease of tissue acquisition allow comprehensive mapping of CE biased domains across the entire ovarian surface at high resolution. We used spatial transcriptomics data generated with 10x Genomics Visium HD on parasagittal sections of XX gonads at E12.5, E14.5, and E16.5 to map CE priming *in situ* at near-cellular resolution (Martinez et al., 2025) (**Fig. 2A**). To resolve spatial organization within the CE, we delineated 4 µm x 4 µm CE bins along the ovarian surface, reconstructed an anteroposterior pseudo-spatial axis, and computed supporting- and SP-identity scores for each bin. At E12.5, CE cells with SP bias were enriched at the extremities of the pseudo-spatial axis near the gonad-mesonephros junction, whereas supporting-biased CE cells were concentrated in the central region (**Fig. 2B** and **C**). This regionalization persisted at E14.5 but became less distinct by E16.5 as the stromal signature declined (**Fig. 2C**). Expression patterns of individual *Runx1* supporting marker and *Nr2f2* SP marker confirmed this organization (**Fig. 2D**). Heatmap visualization of genes differentially expressed in supporting (n = 300) versus stromal (n = 243) cells further confirmed this spatial segregation in the CE (**Fig. 2E, Fig. S2** and **Data S1).** In particular, stromal-biased CE cells at the extremities of the pseudo-spatial axis expressed *Tcf21*, *Pdgfra,* and *Wnt5a*, whereas supporting-biased CE cells in the central region expressed *Wnt6*, *Bmp2*, *Irx3*, *Lgr5*, and *Fst* (**Fig. S2)**. Thus, the molecular signatures that define SP- and supporting-biased CE states in scRNA-seq map onto spatially segregated domains along the ovarian surface.

**Figure 2.**
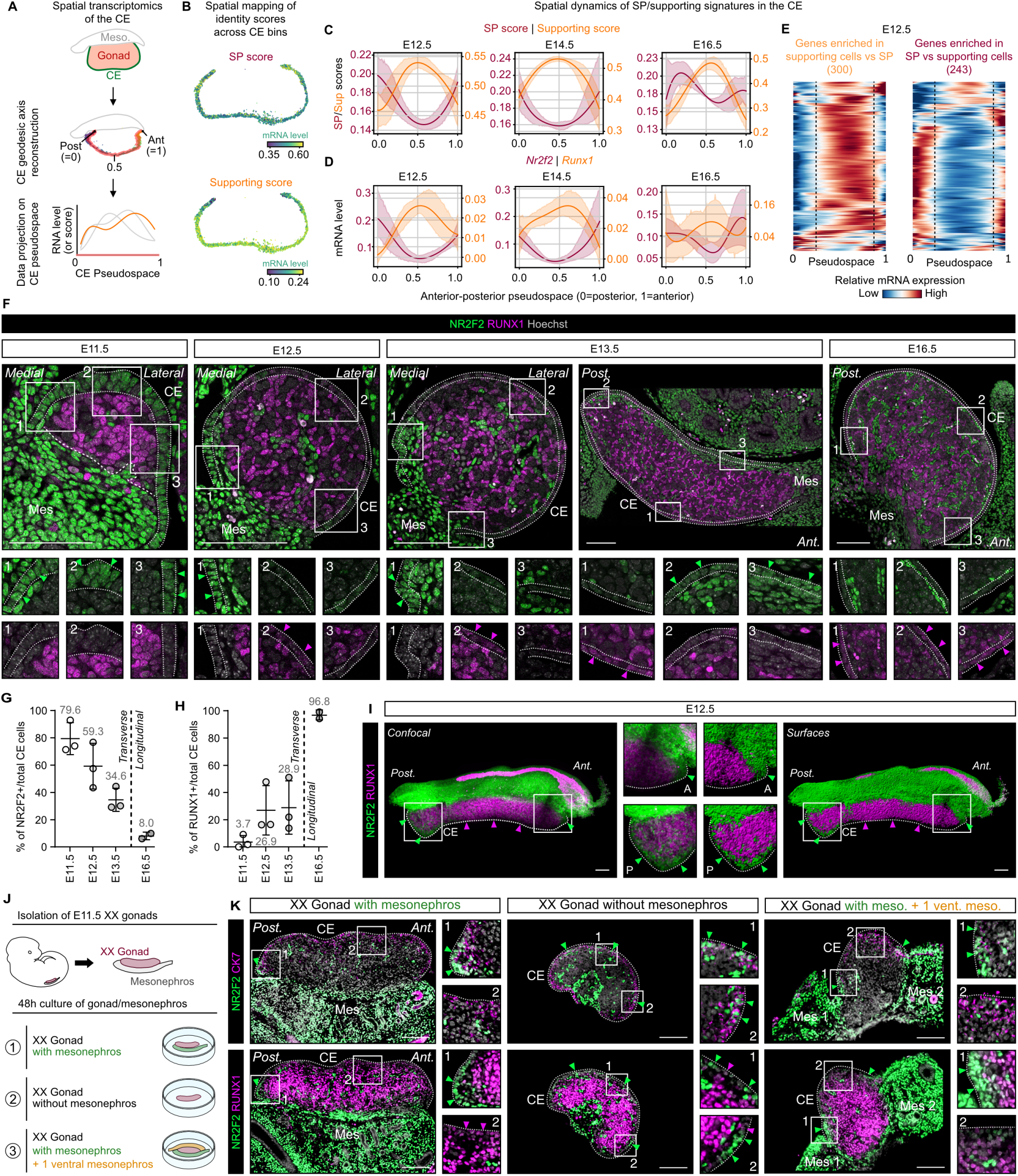
Dynamic spatial patterning of the coelomic epithelium during ovarian development. (A) Spatial transcriptomic experimental design using 10x Visium HD technology on parasagittal ovaries/mesonephros sections. Gonads and CE were annotated, CE paths reconstructed and gene expression/score computed along a pseudospace. (B) Example of stromal and supporting identity score projected along the CE pseudo spatial axis of a developing ovary at E12.5. (C) Lineplots of smoothed stromal- and supporting-identity scores along CE pseudospace at E12.5, E14.5 and E16.5 (Mean with 95% confidence intervals). In the pseudospace, the posterior pole of the ovary is set at 0 and the anterior pole at 1. (D) Lineplots of smoothed *Nr2f2* and *Runx1* expression along CE pseudospace at E12.5, E14.5 and E16.5 (Mean with 95% confidence intervals). Posterior = 0; anterior = 1. (E) Heatmap of gene expression along CE pseudo-space at E12.5 for differentially expressed genes between supporting-cells (pre-Gran, Gran) (n=300) and stromal progenitors (n=243) from the scRNA-seq atlas (log2FC > 1, adj. *p* < 0.01). Genes are ordered by pattern similarity. (F) Immunofluorescence for RUNX1 (supporting) and NR2F2 (SP) in developing ovaries at E11.5, E12.5, E13.5 and E16.5. Dotted lines outline the CE; dashed line marks the boundary between the gonadal and mesonephric compartments. (G and H) Quantification of NR2F2 cells (G) or RUNX1^+^ cells (H) as a percentage of the CE cells at E11.5, E12.5 and E13.5 on transverse sections, and at E16.5 on longitudinal sections. (I).Maximum intensity projection (left) and 3D model generated by isosurface segmentation (right) from whole-mount 3D imaging of E12.5 ovaries stained for NR2F2 and RUNX1. (J) Experimental design of E11.5 XX gonad-mesonephros explant culture. Gonads were cultured for 48h either with the endogenous mesonephros attached (control), after surgical removal of the mesonephros, or with an additional mesonephros placed along the ventral side of the gonad to generate a one-gonad/two-mesonephroi configuration. (K) Co-immunofluorescence for RUNX1 (supporting), NR2F2 (SP), and cytokeratin 7 (CK7, epithelial cells) in E11.5 ovaries +/- mesonephros after 2 days of culture. Scale bars, 100μm.

To validate and refine these spatial domains at cellular resolution, we performed immunofluorescence (IF) on mouse XX gonads from E11.5 to E16.5 using the supporting marker RUNX1 and the stromal/steroidogenic marker NR2F2 (**Fig. 2F**). At E11.5, NR2F2 was broadly expressed across the CE, whereas RUNX1 was not yet detectable, consistent with an initial widespread stromal competence within the CE prior to spatial restriction. By E12.5 and E13.5, IF confirmed the emergence of distinct spatial domains of supporting- and SP-biased CE cells along the ovarian surface. On transverse sections, NR2F2 CE cells localized near the gonad-mesonephros junction, whereas RUNX1^+^ CE cells were restricted to the central region. These regions also exhibited distinct epithelial morphologies: the NR2F2^+^ domain appeared pseudostratified and columnar, while in the RUNX1^+^ domain cells were cuboidal. Over time, the proportion of NR2F2 CE cells progressively declined, while RUNX1 CE cells expanded and eventually covered the entire ovarian surface by E16.5, at which point NR2F2 was almost undetectable in the CE (**Fig. 2G** and **H**).

Whole-mount 3D imaging of cleared E12.5 ovaries further revealed that NR2F2 CE domains form caps at the anterior and posterior poles of the gonad, near the mesonephros, whereas the central CE region was predominantly RUNX1 (**Fig. 2I**). These observations suggest that mesonephric proximity provides positional cues promoting SP bias and/or preventing supporting specification. To test the contribution of the mesonephros to this spatial patterning, we cultured E11.5 XX gonads for 48 hours under three conditions: with the endogenous mesonephros attached, after surgical removal of the mesonephros, or with an additional mesonephros juxtaposed ventrally to generate a one-gonad/two-mesonephroi complex (**Fig. 2J**). Under control conditions with a single attached mesonephros, the *ex vivo* pattern after 48 hours recapitulated the *in vivo* situation at E13.5, with NR2F2 CE domains near the mesonephros and a central RUNX1 CE domain (**Fig. 2K**). Upon mesonephros removal, NR2F2 stromal-biased CE cells no longer formed continuous polar domains but instead appeared as scattered clusters across the ovarian surface, indicating loss of organized stromal-biased caps. Conversely, the presence of an additional mesonephros promoted the formation and expansion of NR2F2 CE domains adjacent to each mesonephros and distorted the central RUNX1 supporting-biased region.

Together, these findings demonstrate that CE fate bias is spatially encoded along the ovarian surface, with SP-biased CE enriched near the gonad-mesonephros interface and supporting-biased CE concentrated centrally. The dependence of NR2F2 stromal-biased domains on mesonephric proximity suggests that mesonephric cues contribute to establish and/or maintain these spatially restricted CE subpopulations.

### Fate-primed ovarian surface epithelium organizes intragonadal domains

As the developing ovarian surface epithelium is partitioned into SP-biased and supporting-biased domains, we next investigated whether these surface compartments seed intragonadal somatic territories. We performed combined *in situ* hybridization (ISH) and IF for the CE marker *Upk3b*, the stromal/steroidogenic NR2F2 and the supporting lineage marker *Bmp2* on XX gonads from E11.5 to E16.5 (**Fig. 3A**). *Upk3b* expression, which is specific to the CE, was used to delineate the surface epithelium from the underlying intragonadal territories. We took advantage of the transient expression of *Bmp2* as a proxy for supporting lineage commitment, as *Bmp2* is expressed in early supporting cells from E11.5 to E13.5 and subsequently downregulated (**Fig. 1N** and **S1F**). At E11.5, NR2F2 was broadly expressed across the CE, whereas *Bmp2* expression was restricted to the central CE domain, indicating that positional bias emerges within the CE before granulosa cell differentiation. At E12.5 and E13.5, *Bmp2* expression remained confined to the central CE, similarly to RUNX1 (**Fig. 2F**). By E16.5, although overall *Bmp2* levels decreased, its expression extended across most of the CE surface. At E12.5 and E13.5, NR2F2 SPs aggregated beneath NR2F2 SP-biased CE, while *Bmp2* pre-granulosa cells localized preferentially beneath the central *Bmp2* CE domain. By E16.5, as the stromal-biased CE domain regressed, the underlying SP-enriched regions became less sharply demarcated, whereas the supporting-enriched intragonadal domain extended beneath the surface epithelium.

**Figure 3.**
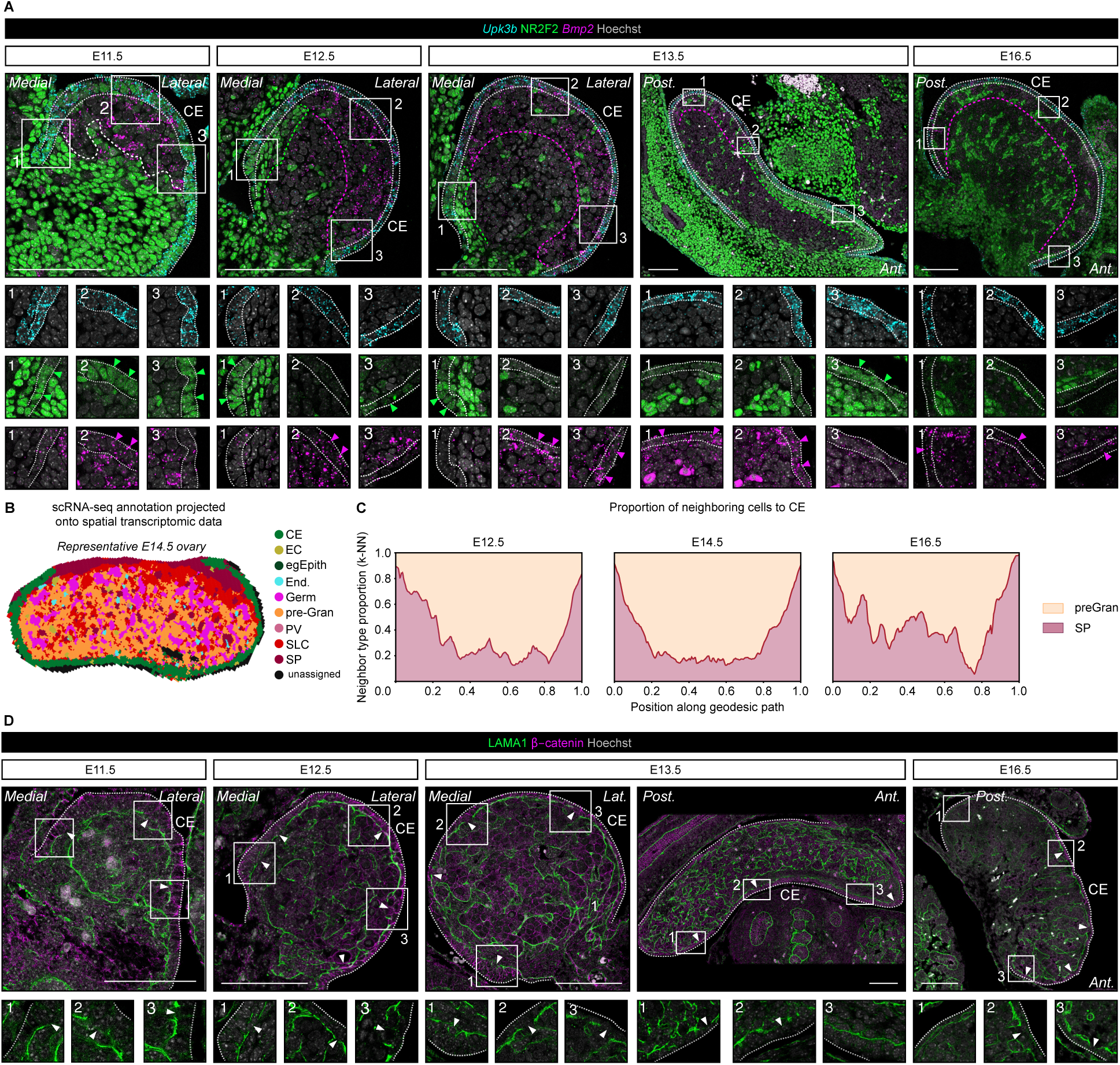
Fate-primed ovarian surface epithelium organizes intragonadal domains. (A) Combined ISH/IF for *Upk3b* (CE/SE), *Bmp2* (supporting) and NR2F2 (stromal progenitor, SP) on developing ovaries at E11.5, E12.5, E13.5 and E16.5. White dotted lines: CE, white dashed lines: boundary between the gonad and the mesonephros, green dashe lines: intragonadal region enriched in NR2F2^+^ cells; magenta dashed lines: intragonadal region enriched in *Bmp2*^+^ cells. (B) Spatial representation of the 4µm bins of a representative E14.5 gonad colored by cell-type signature. (C) Stacked-area plots showing the proportion of intragonadal neighbors (pre-granulosa, Stromal Progenitors (SP)) among the k=100 nearest spatial neighbors (k-NN) of CE bins along pseudospace. (D) Co-IF for β-catenin and LAMA1 at E11.5, E12.5, E13.5 and E16.5. Arrowheads indicate basal lamina discontinuities. Scale bars, 100μm. Abbreviations: CE, coelomic epithelium; EC, erythrocytes; egEpith, extra-gonadal epithelium; End, endothelium; Germ, germ cells; pre-Gran, pre-granulosa cells; PV, perivascular cells; SLC, supporting-like cells; SP, stromal progenitors.

To refine these spatial relationships, we projected scRNA-seq annotations onto the Visium HD spatial dataset to label intragonadal bins (**Fig. 3B**). To evaluate the spatial correspondences between intragonadal cell types and biased-CE progenitors along the ovarian surface, we quantified the relative proportion of SPs and pre-granulosa cells along the CE pseudo-spatial axis (**Fig. 3C**). At E12.5, regions directly beneath SP-biased CE at the ovarian extremities were enriched for SPs, whereas regions underlying the central supporting-biased CE domain contained higher proportions of supporting cells. This complementary pattern persisted at E14.5, with SP-enriched territories remaining aligned with the anterior and posterior caps, and supporting-lineage territories occupying the central ovarian region. By E16.5, as the SP-biased CE domain regressed and the supporting-biased CE domain expanded, the intragonadal SP and supporting territories were less clearly regionalized. Spatial transcriptomics confirmed that intragonadal distributions of pre-granulosa cells and SPs align with the spatiotemporal organization of CE-biased progenitors. To further characterize how biased CE cells can contribute to their corresponding intragonadal compartments, we examined LAMA1 expression to evaluate the integrity of the basal lamina underlying the CE domains (**Fig. 3D**). LAMA1 staining revealed focal discontinuities in the basal lamina beneath both SP-biased and supporting-biased CE domains, at their interface with the corresponding intragonadal SP and pre-granulosa territories. These focal breaks are consistent with localized epithelial-to-mesenchymal transitions that enable fate-biased CE cells to delaminate into the gonad.

Together, these observations support a model in which spatially patterned, fate-primed CE domains contribute to corresponding intragonadal somatic territories, coupling ovarian surface organization to the establishment of supporting and stromal/steroidogenic lineages.

### Early *Wnt5a*+ coelomic epithelium is the major source of granulosa cells

Transcriptional and spatial analyses suggest that pre-patterned CE domains seed both pre-granulosa cells and SPs. However, the spatiotemporal parameters governing their emergence and differentiation remain poorly defined, including the relative contributions of CE progenitors to medullary versus cortical pre-granulosa populations, and whether fetal SPs ultimately differentiate into steroidogenic stromal and thecal cells. To directly test these lineage relationships, we performed inducible *in vivo* lineage tracing using *Wnt5a-rtTA;TetO-Cre;Rosa-tdTomato* mice (hereafter *Wnt5a*^lin^; (Ademi et al., 2022)). Single-cell RNA-seq analyses revealed that at E11.5, *Wnt5a* is expressed in the CE and early SPs but not in pre-supporting and pre-granulosa cells, whereas by E16.5 its expression is restricted to late SPs (**Fig. 4A**). This temporal shift in *Wnt5a* expression enabled two complementary induction windows in *Wnt5a*^lin^ mice: an early window (E11.5-E12.5) to label CE and a late window (E15.5-E16.5) to selectively label fetal SPs (**Fig. 4B** and **C**).

**Figure 4.**
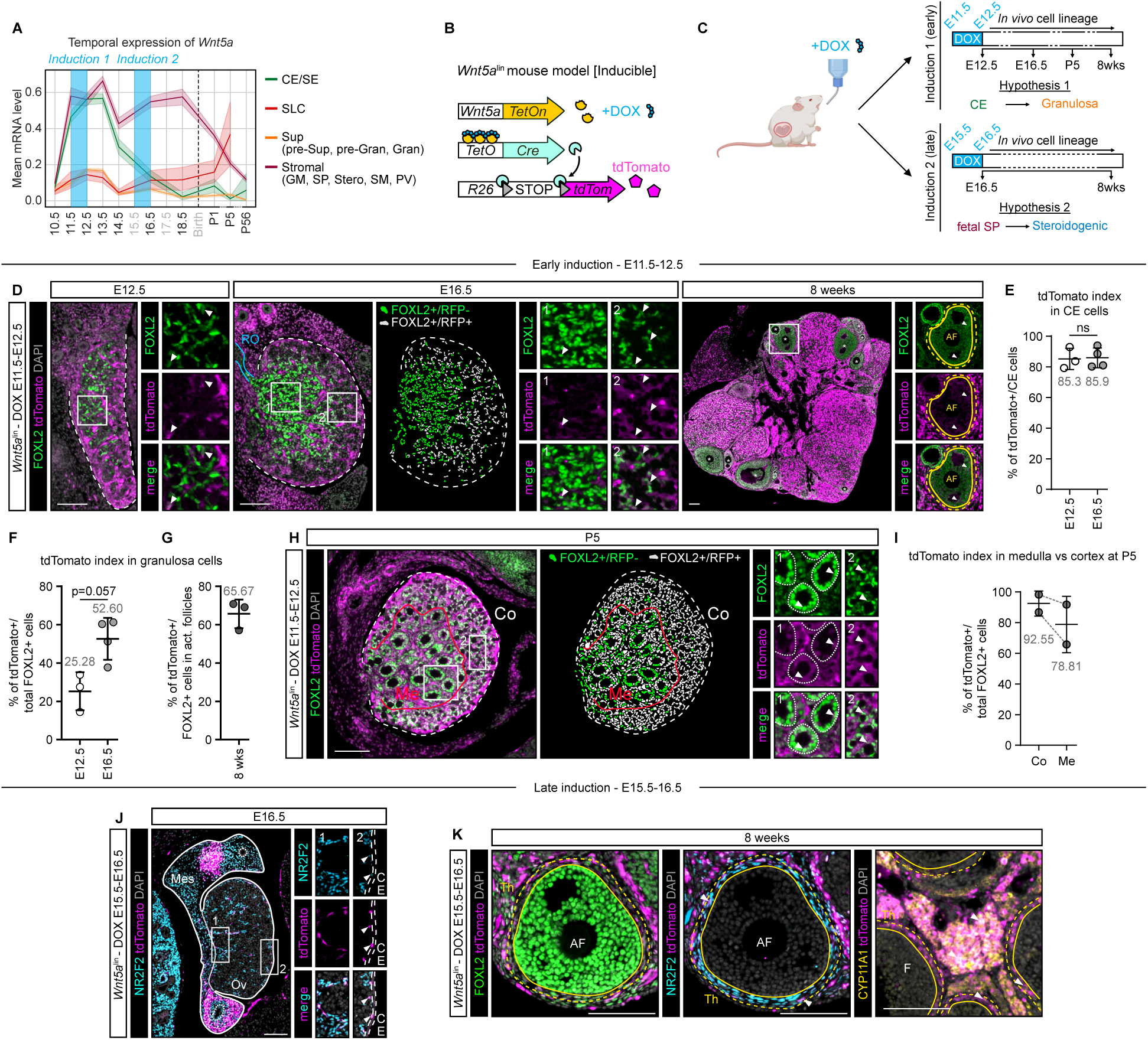
Stage-specific *Wnt5a* lineage tracing reveals CE-derived granulosa cells and fetal SP-derived steroidogenic cells. (A) Mean *Wnt5a* expression across mouse ovarian somatic compartments (CE/SE, SLC, Sup., Stromal) from embryonic day (E)10.5 to post-natal day (P)56. (B) Schematic of the inducible *Wnt5a*^lin^ mouse model (*Wnt5-rtTA;TetO-Cre;R26-tdTomato*). Upon doxycycline (DOX) exposure, Tet-On3G induces Cre expression, excising the STOP cassette in the R26-LSL-*tdTomato* locus and resulting in permanent tdTomato labeling of *Wnt5a*^+^ cells. (C) Experimental design for *in vivo* lineage-tracing. Pregnant females were either exposed to DOX between E11.5 and E12.5 (early induction) and ovaries were collected at E12.5, E16.5, P5, and 8 weeks or exposed to DOX between E15.5 and E16.5 (late induction) and ovaries were collected at E16.5 and 8 weeks. (D) Co-immunofluorescence (co-IF) for tdTomato and FOXL2 in *Wnt5a*^lin^ ovaries at E12.5 and E16.5. (E) Quantification of tdTomato as a percentage of CE cells at E12.5 and E16.5. (F and G) Quantification of tdTomato as a percentage of total FOXL2 pre-granulosa cells at E12.5 and E16.5 (F), and as a percentage of total FOXL2 pre-granulosa in growing follicles at 8 weeks (G). (H) Co-IF for tdTomato and FOXL2 in *Wnt5a*^lin^ ovaries at P5. (I) Quantification of tdTomato cells as a percentage of total FOXL2 granulosa cells in the ovarian medulla (Me) and cortex (Co) at P5. (J) Co-IF for tdTomato and NR2F2 in *Wnt5a*^lin^ ovaries at E16.5. (K) Co-IF for tdTomato with FOXL2, NR2F2 (stromal cells), or CYP11A1 (steroidogenic cells) in *Wnt5a*^lin^ ovaries at 8 weeks. White lines delineate the CE, red lines delineate the ovarian medulla, yellow lines delineate follicular granulosa cells of the antral follicles (AF), yellow dashed lines delineate the thecal compartment. Arrowheads indicate tdTomato /FOXL2 double-positive cells. Dashed lines delineate the CE/SE. Scale bars, 100 μm. Error bars represent mean ± SD. Abbreviations: CE, coelomic epithelium; SE, surface epithelium; SLC, supporting-like cells; pre-Sup, pre-supporting cells; pre-Gran, pre-granulosa cells; Gran, granulosa cells; GM, gonadal mesenchyme; SP, stromal progenitors; Stero, steroidogenic cells; SM, smooth muscle cells; PV, perivascular cells.

Following doxycycline induction at E11.5-E12.5, tdTomato labeled 85% of the CE at E12.5, indicating efficient targeting of the CE (**Fig. 4D** and **E**). Quantification revealed that 25% of FOXL2 pre-granulosa cells were tdTomato^+^ at E12.5, suggesting an early CE contribution (**Fig. 4F**). By E16.5, this proportion increased to 52%, predominantly in the cortex, and in adult ovaries, 65% of FOXL2 granulosa cells were tdTomato (**Fig. 4D, F** and **G**). To distinguish the contribution of CE cells to first versus second waves of folliculogenesis, we analyzed ovaries at P5 after primordial follicle activation (**Fig 4H** and **I**). The first wave of folliculogenesis is initiated in the medulla, whereas subsequent waves arise from a cortical follicle pool. At P5, tdTomato granulosa cells were preferentially localized in the cortex and accounted for 92% of tdTomato /FOXL2 cells, consistent with a predominant CE origin for cortical granulosa cells of the second follicular wave (**Fig 4I**). Notably, a substantial contribution was also observed in the medulla, where 79% of FOXL2 granulosa cells were tdTomato . Together, these data show that early *Wnt5a* CE cells give rise to a major proportion of the granulosa lineage, particularly within the cortical ovary, which harbors the follicle reserve for the reproductive lifespan.

### Late fetal *Wnt5a*^+^ stromal progenitors give rise to steroidogenic stromal and theca cells

We next leveraged the restriction of *Wnt5a* expression to trace the fate of fetal SP. Doxycycline induction at E15.5-E16.5 selectively labeled NR2F2 fetal SPs, with no tdTomato detected in CE or FOXL2 pre-granulosa cells, confirming induction specificity (**Fig. 4J**). In adult ovaries, tdTomato cells were confined to the stromal compartment, largely composed of NR2F2^+^ cells, and excluded from the FOXL2^+^ follicular compartment. Moreover, *Wnt5a*^+^ fetal SPs contribute to CYP11A1 steroidogenic stromal and thecal cells (**Fig. 4K**).

Together, these stage-specific *Wnt5a* lineage-tracing experiments functionally disentangle CE and stromal progenitor fates: early CE cells give rise to the majority of cortical and medullar granulosa cells, consistent with the supporting-biased domains identified at the ovarian surface, whereas late fetal SPs persist as long-lived adult SPs and generate stromal and theca steroidogenic populations.

### Supporting-like cells provide an additional source of granulosa cells

Cross-species transcriptomic analyses predicted that, in addition to CE-derived granulosa cells, SLCs are an additional source of pre-granulosa cells. In the human atlas, SLCs were preferentially connected to preGran-I of the first follicular wave (**Fig. 1A, B** and **E**). Similarly, in the mouse dataset, SLCs connect to *Foxl2*^+^ pre-granulosa cells (**Fig. 1C, D** and **F**), indicating a potential conserved contribution of SLC to the supporting lineage. *Pax8* expression is restricted to SLCs and the rete ovarii, with no expression in CE or pre-granulosa cells (**Fig. 5A** and **B**). This pattern enables specific lineage tracing using *Pax8-Cre;Rosa-tdTomato* mice (hereafter *Pax8^lin^*) (Mayere et al., 2022), allowing us to assess the contribution of *Pax8* SLCs to granulosa cell formation (**Fig. 5C, D** and **Fig. S3**).

**Figure 5.**
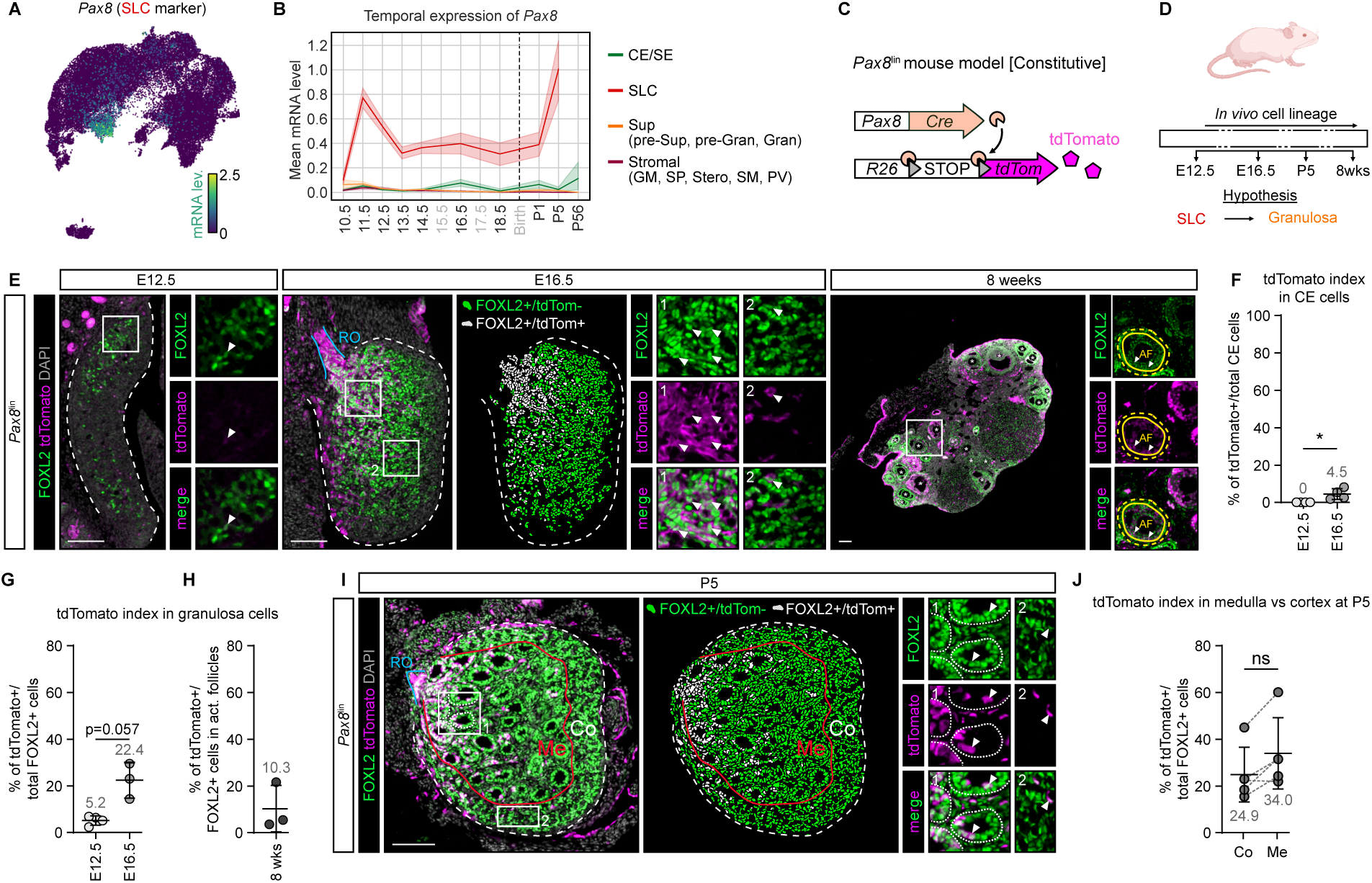
*Pax8□* supporting-like cells represent an additional source of granulosa cells. (A) UMAP representation showing *Pax8* expression in ovarian somatic cells. (B) Mean *Pax8* expression across mouse ovarian somatic compartments (CE/SE, SLC, Sup., Stromal) from embryonic day (E)10.5 to post-natal day (P)56. (C) Schematic of the constitutive *Pax8*^lin^ mouse model (*Pax8:Cre;R26-tdTomato*). *Pax8:Cre* knock-in drives *Cre* recombinase expression under the control of *Pax8* endogenous promoter. Cre-mediated excision of the STOP cassette at the *R26-LSL-tdTomato* locus permanently labeled *Pax8*^+^ cells and their derived cells with tdTomato. (D) Experimental design for *in vivo* lineage-tracing. Ovaries were collected at E12.5, E16.5, P5, and 8 weeks. (E) Co-immunofluorescence (co-IF) for tdTomato and FOXL2 in *Pax8*^lin^ ovaries at E12.5, E16.5 and 8 weeks. (F) Quantification of tdTomato□cells as a percentage of the CE cells at E12.5 and E16.5. (G and H) Quantification of tdTomato□ cells as a percentage of total FOXL2□ pre-granulosa cells at E12.5 and E16.5 (G), and as a percentage of total FOXL2 pre-granulosa in growing follicles at 8 weeks (H). (I) Co-IF for tdTomato and FOXL2 in *Pax8*^lin^ ovaries at P5. (J) Quantification of tdTomato□ cells as a percentage of total FOXL2 granulosa cells in the ovarian medulla (Me) and cortex (Co) at P5. Arrowheads indicate tdTomato□ /FOXL2□ double-positive cells. White lines delineate the CE. Scale bars, 100 μm. Error bars represent mean ± SD. Abbreviations: CE, coelomic epithelium; SE, surface epithelium; SLC, supporting-like cells; pre-Sup, pre-supporting cells; pre-Gran, pre-granulosa cells; Gran, granulosa cells; GM, gonadal mesenchyme; SP, stromal progenitors; Stero, steroidogenic cells; SM, smooth muscle cells; PV, perivascular cells.

At E12.5, tdTomato cells were confined to the SLC/rete region at the hilum and were absent from the CE (**Fig. 5E** and **F**). Notably, ∼5% of FOXL2 pre-granulosa cells were tdTomato^+^, suggesting an early contribution of SLC to granulosa cells (**Fig. 5E** and **G**). By E16.5, this proportion increased to ∼22%, with tdTomato^+^/FOXL2 pre-granulosa cells enriched near the rete ovarii and hilum. By adulthood, ∼10% of FOXL2 granulosa cells were tdTomato, consistent with a decline of the SLC-derived population (**Fig. 5H**). Having established the temporal dynamics of the SLC contribution to granulosa cells, we examined how these SLC-derived granulosa cells are distributed across medullary and cortical compartments at P5 (**Fig. 5I**). At this stage, SLC-derived granulosa cells were present in both medullary and cortical regions in comparable proportions of FOXL2^+^ cells (**Fig. 5J**). A gradient of tdTomato /FOXL2 granulosa cells extended from the hilum toward the center of the ovary, supporting a spatially organized contribution of SLC-derived cells.

Together with the *Wnt5a* lineage-tracing experiments, these findings demonstrate that granulosa cells derived from two spatially distinct progenitor pools: early *Wnt5a* CE progenitors generate the majority of cortical granulosa cells that form the long-lived follicle reserve, whereas SLCs represent an additional source of granulosa cells with a more limited contribution to the adult ovary. This dual origin, supported by both mouse and human scRNA-seq data, refine the classical model of an exclusively CE-derived granulosa lineage.

## Discussion

During early gonadogenesis, proliferating CE cells generate most somatic cells in both XX and XY gonads. Classical dye labeling, lineage tracing, and single-cell transcriptomic studies have established CE progenitors as multipotent, giving rise to both supporting and steroidogenic lineages (Ademi et al., 2022; DeFalco et al., 2011; Karl and Capel, 1998; Liu et al., 2016; Mayere et al., 2022; Mork et al., 2012; Schmahl et al., 2003; Stevant and Nef, 2018, 2019). Yet, the mechanisms that establish these divergent fates, particularly the specification of steroidogenic and supporting progenitors, and the spatiotemporal parameters governing their emergence and differentiation, remain unclear and poorly understood. Recent work on adrenogonadal primordium separation revealed that CE cells are specified toward adrenal or gonadal fates before ingression (Neirijnck et al., 2023), suggesting that specification within the CE may precede integration into the gonad through spatial pre-patterning of progenitor identity.

### Conserved and dynamic coelomic epithelium priming in human and mouse ovarian development

Although ovarian development in humans unfolds over a longer timescale, weeks rather than days as in mice, the dynamics of somatic lineage specification are remarkably conserved. Transcriptomic connectivity between somatic populations, whether assessed globally or across developmental stages, shows similar patterns in human and mouse, as do cell-type proportions and proliferation dynamics. The only difference concerns the early SP population, which in humans appears to separate early pre-granulosa cells from the CE. It remains unclear whether this transitional state is absent in mice or instead represents a transient population that is too short-lived to be captured. Our study also identifies transcriptionally and spatially distinct fate-biased CE domains with progenitor subpopulations overlying the developing ovary. CE cells co-expressing either SP (*Arx/ARX, Pdgfra/PDGFRA, Tcf21/TCF21*) or supporting (*Bmp2, Irx3/IRX3, Runx1, Wnt6/WNT6, LHX2*) markers exhibit transcriptomic signatures resembling their respective target lineages, consistent with *bona fide* priming. At early stages, in both mouse (E11.5) and human (PCW5–6), most CE cells are multipotent and NR2F2 . In mice, a subset of these cells already expressing *Bmp2*, marking the onset of positional bias. We propose that a first wave of supporting cells is generated before or around E11.5 through progressive downregulation of progenitor markers (e.g., NR2F2, TCF21) and acquisition of supporting identity upon entry into the gonad. From E12.5 onward, CE cells become spatially organized: supporting-primed (*Upk3b /Bmp2*) cells expand across the ovarian surface, whereas steroidogenic-primed (*Upk3b* /NR2F2) cells remain restricted near the mesonephros. At these stages (E12.5-E14.5), supporting-primed CE progenitors generate additional supporting cells, while steroidogenic-primed cells give rise to SPs. By E16.5, CE cells are almost exclusively supporting-primed and contribute to pre-granulosa and surface epithelial cells.

Spatial transcriptomics and immunofluorescence analyses revealed that intraovarian SP and pre-granulosa cells are positioned directly beneath their corresponding CE subdomains, consistent with local delamination of primed progenitors seeding spatially distinct intragonadal niches. These findings establish a direct, spatially organized contribution of CE subpopulations to ovarian lineage formation. In humans, CE progenitors display comparable dynamic priming but over a longer timescale. The sequence progresses from an early steroidogenic bias to a transitional coexistence of both progenitor types, followed by a predominance of supporting-biased CE cells as ovarian development advances.

Our observations align with work by Bunce et al. (Bunce et al., 2023), who reported center-biased ingression of supporting progenitors in both XX and XY gonads, producing higher supporting cell density in the central domain. Using a *Runx1-EGFP* reporter, they demonstrated greater accumulation of supporting precursors in the gonadal center. Consistently, our NR2F2/*Bmp2* combined *in situ* hybridization (ISH) and IF at E11.5 revealed that while most CE cells are NR2F2, faint *Bmp2* /NR2F2^+^ CE cells are detectable in the central ovarian region. Together, these results support a model in which regional patterning of CE progenitors determines lineage allocation.

### Mesonephros as a positional regulator of coelomic epithelial fate

In mice, from E12.5 onward, supporting-biased CE cells expand broadly across the ovarian surface, whereas stromal/steroidogenic-biased CE cells remain confined near the mesonephros. This spatial segregation suggests that mesonephric proximity provides positional cues that favor stromal/steroidogenic bias and/or prevent supporting bias specification when both CE subpopulations coexist. Consistent with this hypothesis, *ex vivo* culture experiments show that CE cells cultured adjacent to the mesonephros display enhanced NR2F2 expression, indicating that local signals either promote steroidogenic identity and/or inhibit supporting fate specification. We propose a model in which mesonephros-gonad crosstalk establishes a spatial gradient that sustains SP priming in CE progenitors located near the mesonephros. These results suggest that local environmental signals play a critical role in the specification of somatic lineages, and that they should be taken into account in protocols designed to generate functional ovarian organoids from iPS cells. Future work integrating ligand-receptor inference with targeted perturbations will be crucial to identify factor(s) or signaling pathways mediating this interaction and to determine how these mechanisms influence ovarian development and pathophysiology.

### Conserved sequence but sex-specific timing of steroidogenic lineage differentiation

In mammals, testicular and ovarian differentiation follow distinct temporal trajectories, with testis differentiating earlier and more rapidly (Nicol et al., 2022; Stevant and Nef, 2019; Wilhelm et al., 2013). This divergence is most evident in androgen-producing cells: fetal Leydig cells begin secreting androgens shortly after testis differentiation (E12.5 in mouse; 6-7 PCW in human), whereas ovarian theca cells emerge only late in fetal life and do not acquire full steroidogenic activity until after birth, most prominently at puberty, when they become LH-responsive during folliculogenesis (Barsoum and Yao, 2010; Fortune and Armstrong, 1977; Magoffin, 2005; Mannan and O’Shaughnessy, 1991; Richards et al., 2018). Our mouse and human data reveal a conserved trajectory in which a subset of CE cells acquires a steroidogenic-primed identity during a restricted developmental window and contributes directly to intraovarian SPs. These SPs proliferate actively and ultimately constitute roughly half of all ovarian cells in both species, mirroring the marked expansion of SPs observed in the developing mouse testis, where they are also predominant at birth (Ademi et al., 2022). In both human and mouse, ovarian SPs express canonical markers (*Arx/ARX, Pdgfra/PDGFRA,* and *Tcf21/TCF21*), which are also characteristic of testicular SPs (Brennan et al., 2003; Cui et al., 2004; Eliveld et al., 2019; Estermann et al., 2025; Ge et al., 2006; Kilcoyne et al., 2014; Miyabayashi et al., 2013; Perea-Gomez et al., 2025; Qin et al., 2008; Shen et al., 2021). Lineage tracing further showed that fetal *Wnt5a* SPs persist and give rise to steroidogenic theca and stromal cells in the ovary as well as FLCs in the testis (Ademi et al., 2022). Thus, although the timing differs, the overall sequence of SP specification expansion, and differentiation is likely to be conserved across sexes and species.

### A dual origin for granulosa cells

Our study challenges the long-standing view that granulosa cells derive exclusively from the CE and instead supports a convergent, dual-origin model in both mouse and human ovaries. By integrating single-cell transcriptomics, spatial mapping and lineage tracing, we show that granulosa cells arise from two transcriptionally and spatially distinct progenitor pools. CE-derived progenitors generate the majority of cortical granulosa cells that contribute to the long-lived follicle reserve, whereas SLCs give rise to an additional granulosa subpopulation with a more limited contribution to the adult granulosa cells. This dual origin adds an important layer of complexity to ovarian lineage architecture and helps reinterpret follicular heterogeneity. The developmental origin of granulosa cells likely influences how distinct granulosa subsets interact with oocytes, proliferate, and respond to paracrine, endocrine and metabolic signals throughout ovarian life.

## Material & Methods

### Single-cell RNA sequencing analysis

#### Human data analysis

Raw count matrices from Garcia-Alonso et al. (Garcia-Alonso et al., 2022) were downloaded from their authors’public archive and human single cell raw counts from Lardenois et al. (Lardenois et al., 2026) were provided directly by the corresponding author. Dimensionality reduction was performed using SCVI with batch correction by sample and processing laboratory or by processing laboratory only. A SCANVI model for cell type prediction was trained using the annotations from both Garcia and Lardenois datasets.

#### Human data annotation

To refine cell type predictions obtained with the SCANVI model, the dataset was subset by developmental time point. For each subset, UMAP embeddings were generated and clustering was performed. Small clusters composed of barcodes with low UMI counts and low numbers of detected genes were removed as low-quality cells. Remaining clusters were annotated based on marker gene expression, SCANVI predictions, and the annotations reported in the original publications.

#### Human data embedding

As correcting batch effect at the sample level can conceal biological relevant information such as developmental stage, we created neighbor graphs using intermediate correction strategies. Specifically, we generated SCVI latent spaces corrected either by sample or by laboratory and built the corresponding neighbor graphs. Graph connections were then interpolated between these two correction schemes to produce 10 intermediate correction steps. After visual inspection, the third interpolation step was selected for downstream visualization, as it preserved developmental stage structure while reducing batch effects. For visualization, gene expression levels were log-normalized (target sum = 10000).

#### Human data lineage analysis and scoring

Lineage relationships within the human somatic compartment were inferred using a modified version of PAGA (Wolf et al., 2019). Conductance-based PAGA (cPAGA) in which the connectivity matrix between groups of cells is measured using intergroup conductance in the neighbor graph. In short, conductance measures the proportion of edges that are connected to neighbors that between two groups over the total number of edges. CE cells Sup and SP scoring was performed by training an ElasticNet regressor on Sup and SP cell categories and predicting the score of CE cells.

#### Mouse data preprocessing

Demultiplexing, alignment, barcode filtering, UMI counting and gene mapping were performed as described in the original publications: Mayère et al., 2022 for E10.5, E11.5, E12.5, E13.5 and E16.5 (Mayere et al., 2022); Niu et Spradling, 2020 for E14.5, E18.5, P1 and P5 (Niu and Spradling, 2020); and Rossitto et al., 2022 for P56 (Rossitto et al., 2022). For each sample, low-quality or empty droplets were removed by filtering barcodes with low UMI counts, following the procedure described in (Mayere et al., 2022). Otherwise, they were removed by visual inspection of the UMI counts density curves. Doublet scores were predicted using Scrublet v0.2.3 (Wolock et al., 2019) via the Scanpy v1.9.8 (Wolf et al., 2018) wrapper. Doublet scores thresholds were defined by identifying the closest local minima to the main peak of scores distribution, and barcodes above this threshold were removed. Additional quality controls and filtering were performed using scanpy.pp.calculate_qc_metrics for mitochondrial and ribosomal content. Cells with >20% mitochondrial counts were excluded. Potential XY-contaminating barcodes were identified based on *Ddx3y* expression (discarded if >0). All samples were then merged into a single AnnData object retaining 68,366 cells and 19,961 genes in total: 6,798 (E10.5), 5,935 (E11.5), 9,230 (E12.5), 10,866 (E13.5), 9,555 (E14.5), 6,223 (E16.5), 3,186 (E18.5), 4,932 (P1), 5,281 (P5), and 6,360 (P56).

#### Dimensionality reduction and batch correction of mouse data

Dimensionality reduction and batch correction were performed with scvi.model.SCVI from scVI-tools v1.3.0 (Gayoso et al., 2022) on raw counts data. The model was trained using the samples IDs as batch keys to correct for the batch effects, with the following hyperparameter: 100 latent_space, 3 hidden layers with 128 nodes each.

#### Mouse data counts normalization

For expression visualization, differential expression, scoring, and model training, counts were log-normalized. Specifically, for each cell, UMI counts for each gene were divided by the total UMI counts of the cell, multiplied by a factor of 10,000, and log-transformed using scanpy.pp.normalize_total and scanpy.pp.log1p respectively. Cell cycle scoring was performed with scanpy.tl.score_genes_cell_cycle using the gene set from (Tirosh et al., 2016) as input.

#### Mouse data clustering, UMAP embedding, and annotation

For the full atlas, neighbors were computed in the SCVI latent space using scanpy.pp.neighbors with default parameters. Clustering was performed with scanpy.tl.leiden at resolution 1. We used PAGA on subsampled (ncells=200) clusters to optimize the global UMAP topology, we generated and plotted the PAGA graph with default parameters (scanpy.tl.paga) and used PAGA positions as seeds for the UMAP embedding. UMAP embedding was computed with scanpy.tl.umap function with min_dist parameter set to 0.2.

scVI based dimensionality reduction and batch correction are highly reliable, at the cost of a substantial loss of temporal information. To account for the time-dependent annotations and avoid inconsistencies, we annotated the dataset at two levels. First, a coarse annotation across the full dataset was performed using known marker expression. Second, refined annotations providing greater accuracy and reducing inconsistencies were carried out at each developmental stage using the coarse annotation as a guide. Each timepoint was clustered and embedded independently using the same preprocessing and clustering pipeline, except that PAGA subsampling was reduced to 100 cells per cluster. Known stage-specific markers were used to annotate clusters. Because temporal subsets contain fewer cells, rare cell populations were sometimes difficult to isolate. In ambiguous cases, subclustering was performed with a lower Leiden resolution (0.15 or 0.25).

Additionally, both E10.5 and E11.5 subsets required a specific annotation strategy due to their lower differentiation state and fraction of gonadal cells. For the E10.5 subset, we retained the coarse annotation and Leiden clustering from the full atlas and defined the mesonephric mesenchyme (MM) population by their low *Gata4* and high *Gata2* expression. For the E11.5 subset, Leiden clustering performed either on the full atlas or on the E11.5 subset did not separate gonadal and extragonadal stromal cells. We therefore isolated all stromal cells (*Tcf21*^+^/*Gata4*^+^/*Gata2*^+^) using this initial clustering. To discriminate gonadal from extragonadal cells, we applied a k-nearest neighbors (kNN)-based co-expression search to identify the ten genes strongly co-expressed with *Gata4* and *Gata2* respectively. After verifying the specificity of these gene sets, we computed expression scores using sc.tl.score_genes, scaled them with a MinMax scaler, and assigned each cell to the gonadal or extragonadal compartment based on the dominant score.

#### Sub-atlases

To focus on the two major somatic lineages of the ovary, we extracted a sub-atlas restricted to ovarian somatic populations. scVI latent space was not recomputed, but the same preprocessing and clustering strategy as in the full atlas was applied. PAGA threshold parameter was set to 0.2, and UMAP min_dist parameter set to 0.3. Additional subsets were generated by isolating ovarian somatic populations at each stage, following the same strategy used for the time-point subsets described above.

#### Scoring

To create SP and supporting scores, we trained an ElasticNet model in a one-versus-rest fashion on CE/SE, SLC, SP and supporting (pre-Gran, Gran) cells from E12.5 to P5. Model parameters were set to an L1 ratio of 0.5, alpha of 0.01, and tolerance of 1e-4. Model parameters were set to keep only positive features, and with an L1 ratio of 0.5, alpha of 0.01, and tolerance of 1e-4. To avoid bias from overrepresented cell types, we randomly subsampled each group to match the size of the smallest population prior to training. By comparing each group to the other cells, this approach attributes to each gene a coefficient to create a transcriptomic signature. Genes with high descriptive power are attributed to high coefficients. Consequently, the accuracy of this method depends heavily on the annotation quality and the cellular composition of the subset. For reliability, E10.5 and E11.5 cells were excluded since lineages are not yet committed, and somatic differentiation has not occurred at this stage. We also restricted the analysis to fetal and perinatal data and focused exclusively on ovarian somatic cells (CE/SE, SLC, supporting lineage and steroidogenic/stromal lineage). All the other cell types (adrenosympathetic, erythrocytes, endothelial, germ, immune, mesonephric tubules, mesonephric mesenchyme, perivascular) were excluded from training set.

#### Differential expression analysis

Differential expression analysis was performed using sc.tl.rank_genes_groups and the Wilcoxon method.

### Spatial transcriptomics analysis

#### Data collection

Detailed methods for generating the full spatial transcriptome dataset of the developing and adult mouse ovary are described in (Martinez et al., 2025). Briefly, ovaries from fetal CD-1 outbred mice were collected at E12.5, E14.5 and E16.5, transferred to cryomolds, embedded in OCT, flash frozen and stored at -80°C. Cryoblocks were sectioned using a Leica cryostat to produce 10µm sections, which were stained with Hematoxylin and Eosin (H&E). H&E-stained slides were imaged at 20x, and the highest quality and most representative slide was selected for sequencing. Slides were processed at the University of Colorado Anschutz Medical Campus Genomics Shared resource (RRID: 021984) following the Visium HD Spatial Gene Expression Reagent Kits User Guide (CG000685 Rev B). Sections were hybridized overnight with 10x Genomics’ whole transcriptome probe pairs. The prepared Visium HD slides were processed on the CytAssist instrument (10x Genomics) for spatial barcoding, followed by library preparation. Libraries were pooled and sequenced on an Illumina NovaSeq X Plus system (150bp x 10bp x 10bp x 150bp), generating approximately 500 million reads per capture area.

#### Bins annotation

Raw sequencing and imaging data from 10x Visium HD were processed with spaceranger (v3.1.2) as described in (Martinez et al., 2025). Resulting Zarr datasets were then processed with the spatialdata python library. Initial bin annotation was performed using a logistic regression model trained on single-cell RNA-seq data restricted to genes also detected in the spatial dataset. To prevent overfitting and reduce bias from over-represented classes, cells were randomly under-sampled to 100 cells per population using RandomUnderSampler. The LogisticRegression model was trained using the following parameters: saga solver, elasticnet regularization (l1_ratio=0.8), a stopping criteria tolerance 0.01, and inverse regularization strength 0.8. The model achieved excellent predictive performance, with AUC values of 1.0 for most cell classes, and 0.99 for CE/SE, SP and SLC populations. Precision and recall were also high, with the lowest average score observed for SLCs (precision 0.85) and MM cells (recall 0.83).

Spatial bins were predicted at 2, 4 and 8µm resolutions using the trained model. Because individual bins can contain transcripts from multiple cell types (e.g. pre-Gran and germ cells), predictions were strengthened using the spatial context of each bin. Specifically, logistic regression prediction scores for each bin were adjusted based on the scores of its eight nearest neighbors. For example, if a bin showed mixed Germ and pre-Gran prediction scores, but its neighbors display strong Germ prediction score, the Germ score for that bin was increased accordingly. This smoothing step was implemented by computing the matrix multiplication between the neighbor connectivities and logistic regression prediction probability matrices. Finally, bins with low or ambiguous prediction confidence across all classes were also left unassigned. The confidence threshold was adapted to each resolution, as smaller bins contain fewer transcripts and therefore have lower prediction scores.

#### Surface epithelium path reconstruction

Gonadal areas and associated bins were manually annotated using Napari. Each gonad was aligned using PCA applied to bins spatial coordinates. Since SLCs are progenitors of the rete cells and are located at the anterior pole of the gonad, the averaged SLC prediction score was used to orient each sample. Only gonadal sections with a visible gonad-mesonephros junction were retained for further analysis.

Bins annotated as coelomic epithelium (CE) were extracted and used to reconstruct a path along the ovarian surface. First, a spatial neighbor graph of CE bins was generated using scikit-learn’s kneighbors_graph. Long connections were pruned to avoid artifacts caused by isolated bins or links between opposite CE poles. The shortest paths between all bin pairs were computed using NetworkX (all_pairs_dijkstra_path_length). Terminal points were defined as the pair of bins separated by the longest geodesic distance, and the shortest path between these endpoints was extracted using networkx.shortest_path. All CE bins were then projected onto this path using a KD-tree approach. For each CE bin, the nearest path bin was identified, and distances were computed as cumulative sums of distances from root bin along the path. For visualization, bins were ordered along the obtained pseudospace. Because spatial transcriptomic data are sparse, expression values were smoothed with a generalized additive model (linearGam function of pyGam package), and visualized as heatmaps or curves using seaborn.

#### Scoring of CE and Gene selection for heatmaps

An alternative scoring model was trained using genes detected in both spatial and single-cell RNA-seq datasets. CE bins were then scored using this model. Heatmaps were generated using differentially expressed genes with a log2 fold-change >1 and a corrected p-value<0.001 between supporting cells and SPs of the scRNAseq atlas. Differential expression was assessed with a rank-sum test (log2FC > 1, adj. *p* < 0.01).

#### Proportion of CE neighbors

For visualization, cell annotations were merged into two lineage categories: Supporting (pre-gran and pre-Sup.) and stromal (gSP, mSP, GM). For each CE bin in each gonad and developmental stage, the proportion of neighboring cells belonging to each lineage was calculated using k = 100 nearest neighbors. Profiles were then interpolated to normalize path length across gonads and averaged per timepoint for plotting.

### Fetal gonad collection and *in vitro* culture

Fetal gonads were collected from C57BL/6 embryos obtained from timed matings (E0.5 = day of vaginal plug). Pregnant females were euthanized by cervical dislocation, embryos were dissected, and gonads were isolated at E11.5. As sex cannot be determined morphologically at this stage, embryos were genotyped by PCR detection of the *Sry* gene using the following primers: FW 5′-GTCAAGCGCCCCATGAATGCAT-3′ and RV 5′-AGTTTGGGTATTTCTCTCTGTG-3′. *Sry*-negative (female) gonads were used for *in vitro* culture.

Gonads were placed on Millicell-CM membrane inserts (0.4 μm) under three conditions: (i) with attached mesonephros, (ii) without mesonephros, or (iii) positioned between two mesonephroi. Explants were cultured for 48 h at 37 °C in 5% CO in DMEM/F12 supplemented with 10% FBS and gentamicin (0.04 mg/mL), with medium changed after 24 h. Tissues were then fixed in 4% PFA for immunofluorescence analyses.

### Wild-type, transgenic mouse models and cell lineage tracing

All animal procedures conformed to institutional and European regulations and were approved by the Service de la Consommation et des Affaires Vétérinaires (SCAV) of the Canton de Genève (licenses number GE67-19, GE214-19, GE379a and GE35). Animals were maintained on a mixed genetic background under a 12-h light/12-h dark cycle. with ad libitum access to food and water.

The alleles used in this study were previously described: *Wnt5atm1(Tet-On 3G)Nef*, *(tetO)7-CMV-Cre (TetO:Cre) Pax8tm1.1(cre)Mbu/J* and *Gt(ROSA)26Sortm9(CAG-tdTomato)Hze* (Ademi et al., 2022; Bouchard et al., 2002; Madisen et al., 2010; Perl et al., 2002). For constitutive tracing of *Pax8*^+^cells, we generated *Pax8*^lin^ mice (*Pax8tm1.1(cre)Mbu/J*^+/ki^;*Gt(ROSA)26Sortm9(CAG-tdTomato)HzeR26RFP/^ki/ki^*). For inducible lineage tracing of *Wnt5a*+ cells, we generated *Wnt5a*^lin^ mice (*Wnt5atm1(Tet-On3G)Nef^+/ki^;(tetO)7-CMV-Cre/^+/ki^;Gt(ROSA)26Sortm9(CAG-tdTomato)HzeR26RFP/^+/ki^*). For *Wnt5a*^lin^, doxycycline (DOX) was delivered in drinking water (pH 6.5) at 2 mg/mL during E11.5-12.5 or E15.5-E16.5. For spatial transcriptome experiments, CD-1 outbred mice were housed in an AALAC-accredited barrier facility at the University of Colorado Anschutz Medical campus. All mice were handled and cared for following accepted National Institute of Health guidelines. All experiments were conducted with the approval of the University of Colorado Anschutz Medical Campus Institutional Animal Care and Use Committee (IACUC protocol # 01262). Embryos were staged by detection of a vaginal plug (E0.5), postnatal stages were assigned by day of birth (P0). Ovaries were collected at fetal and adult time points as indicated to follow the fate of *Pax8+* or *Wnt5a+* cells across development.

### Tissue processing, immunofluorescence and *in situ* hybridization analyses

Embryonic samples were collected at defined stages, fixed in 4% (*w/v*) paraformaldehyde overnight, processed for paraffin embedding, and sectioned into 5 µm sections. Immunofluorescence and Hoechst 33342 (H3570, Invitrogen) staining were performed as described in (Tang et al., 2020) or (Ademi et al., 2022). Primary antibodies included: mouse anti-NR2F2 (PP-H7147-00, R&D, 1/200), rabbit anti-RUNX1 (ab92336, Abcam, 1/500), rabbit anti-LAMA1 (L9303, SIGMA, 1/200), mouse anti-ß-Catenin (BD-610153, BD Biosciences, 1/100), rabbit anti-FOXL2 (1:500, gift from Dagmar Wilhelm, University of Melbourne, Australia), mouse anti-αSMA (1:500, gift from Christine Chaponnier, University of Geneva, Switzerland), rabbit anti-PAX8 (1:500, 10336-1-AP, Proteintech), mouse anti-RFP (DsRed) (1:250, sc-390909, SantaCruz Biotechnology, USA), goat anti-RFP (1:50, 200-101-379, Rockland Immunochemicals Inc., USA), rabbit anti-RFP (1:250, 600-401-379, Rockland Immunochemicals Inc., USA) and rabbit anti-CYP11A1 (1:500, gift from Dagmar Wilhelm, University of Melbourne, Australia). RNAscope *in situ* hybridization was performed using the RNAscope Multiplex Fluorescent Reagent Kit v2 (Advanced Cell Diagnostic) using probes for *Bmp2* (406661-C3) and *Upk3b* (568561) following the manufacturer’s instructions.Fluorescent images were acquired on Axio Imager M2 or Z1 microscope (ZEISS, Germany) equipped with Axiocam 702 mono camera or MRm camera (ZEISS, Germany), and or on a Zeiss LSM 780 confocal microscope with a 40x/1.2 W objective. Images were minimally adjusted for global levels in ZEN (ZEISS), processed in Fiji (Bethesda, MD, USA) and assembled using OMERO (https://www.openmicroscopy.org/omero/) or Affinity.

### Immunofluorescence quantification and statistical analysis

Cell quantifications were performed using QuPath-0.5.1. Nuclei were detected by DAPI intensity, cells were classified as positive/negative based on mean signal intensity thresholds for FOXL2, tdTomato, NR2F2 or RUNX1 (thresholds set per experiment). For each condition, 2-5 biological replicates (whole-ovary sections) were analyzed. Results are reported as the percentage of tdTomato^+^, NR2F2^+^, or RUNX1^+^ cells (Mean ± SD).

Statistical analyses were performed in GraphPad Prism (version 10). Pairwise comparisons used two-tailed Mann-Whitney U tests. Significance was defined as P < 0.05. n biological replicates and statistical details are provided in Figure legends.

### Whole-mount immunofluorescence for confocal imaging

E12.5 ovary/mesonephros complexes were collected and fixed in 4% paraformaldehyde 30 min at room temperature, dehydrated through a methanol/PBS gradient (25%, 50%, 75%, 100% methanol), and stored overnight in 100% methanol at -20°C. Samples were then rehydrated through the reverse methanol/PBS gradient, permeabilized in PBS/0.1% Triton X-100 for 30 min, and blocked for 1 h in PBS/0.1% Triton X-100/10% horse serum. Samples were incubated overnight at 4°C with primary antibodies diluted in blocking solution: mouse anti-NR2F2 (PP-H7147-00, R&D, 1/200) and rabbit anti-RUNX1 (ab92336, Abcam, 1/500). After washes in PBS/0.1% Triton X-100, samples were incubated overnight at 4°C with fluorophore-conjugated secondary antibodies (1:500). Samples were mounted in CUBIC Reagent II containing 2% agarose and imaged on a spinning-disk confocal microscope using a 20x/0.8 objective. Images were minimally adjusted for global levels and processed in Imaris.

## Additional resources

An interactive viewer will be available upon publication for further exploration [https://www.unige.ch/medecine/nef/datasets].

## Acknowledgments

We thank Coralie Gobbo, Karim Hammad and Violaine Regard for technical assistance. We are grateful to the Bioimaging platform and to the animal facility at the Faculty of Medicine, University of Geneva, for their support. We also thank the members of the Nef laboratory for helpful discussions.

## Funding

Swiss National Science Foundation grant 31003A_173070 (SN)

Swiss National Science Foundation grant 310030_200316 (SN)

Swiss National Science Foundation grant 10.001.251 (SN)

Département de l’Instruction Publique of the State of Geneva (SN)

Agence Nationale de la Recherche grant ANR-23-CE14-0012 HeteroSex (MCC)

Agence Nationale de la Recherche grant ANR-24-CE14-2476 Anaform (MCC)

CU Anschutz Medical Campus, Department of Pediatrics (JM)

National Institutes of Health grant #R00HD103778 (JM)

Agence Nationale de la Recherche grant ANR-22-CE17-0059 (GL)

Electricité de France-V3.203 (GL)

## Author Contributions (CRediT)

Conceptualization: CD, CM, S N; data curation: CD, CM, MG; formal analysis: CD, CM, MG; Investigation: CD, CM, AG, LB, PB, HA, AR, FK, CG; methodology: CD, CM, AG, LB; visualization: CD, CM, MG, AG; resources: AM, TG, JM, MCC, GL; funding acquisition: SN, MCC, JM, GL; supervision: SN; writing - original draft: CD, SN; writing - review & editing: CD, CM, MG, AG, SN, MCC, JM, DW.

## Competing Interests

The authors declare no competing interests.

## Data and materials availability

All data needed to evaluate the paper are present in the paper and in the Supplementary Materials. The scRNAseq datasets are available at NCBI Gene Expression Omnibus GEO: GSE184708.

## Abbreviation

CE/SE: coelomic epithelium
SP: stromal progenitors
A: anterior pole
P: posterior pole
Mes: mesonephros.

---

**Supplementary Figure S1.**
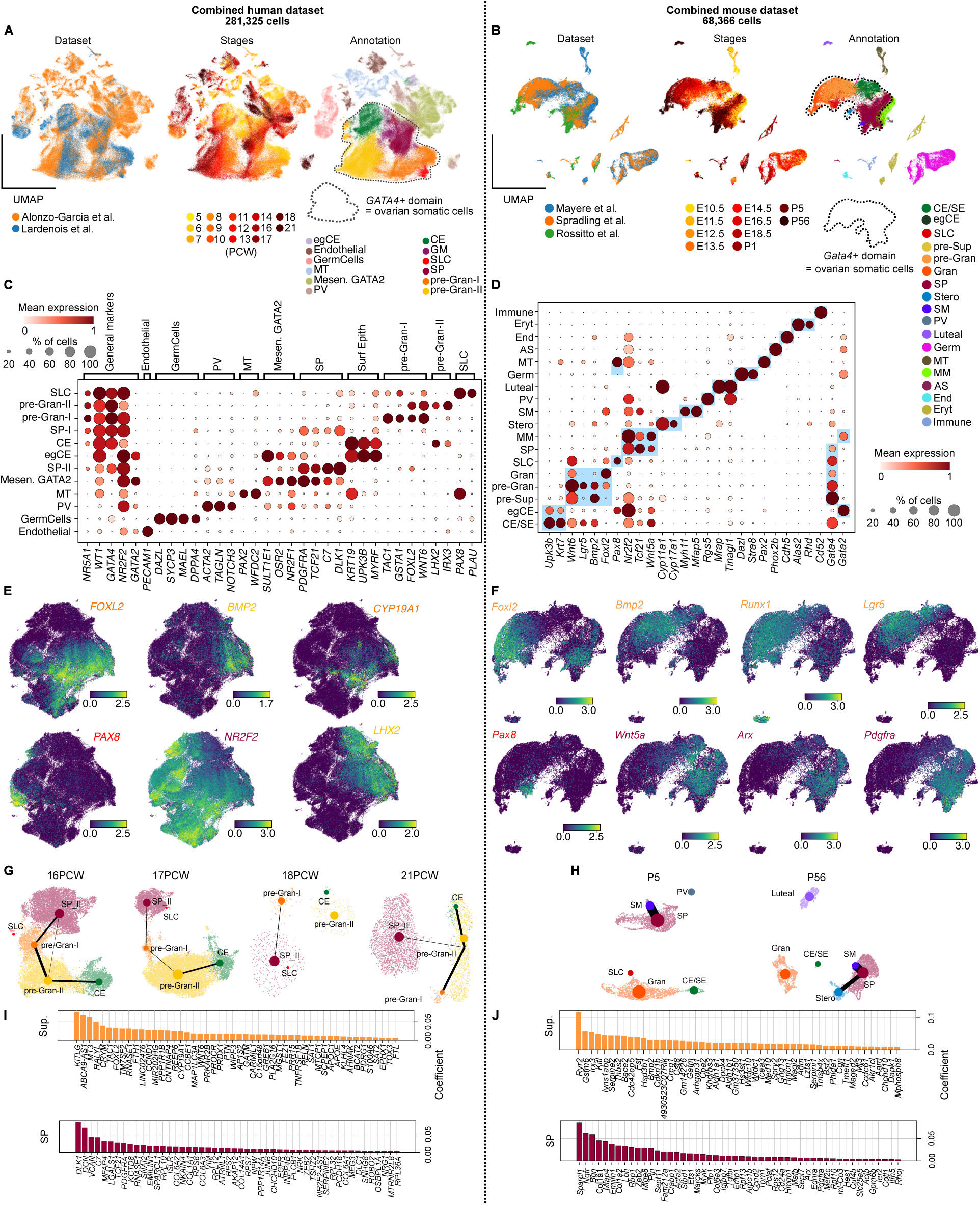
related to Figure 1. (A and B) UMAP representations of single cells from human (n = 281,325) (A) and mouse (n = 68,366) (B) datasets. Cells are colored by dataset, developmental stage (post-conceptional week (PCW) in human, embryonic day (E) and post-natal day (P) in mouse and cell type annotation. (C and D) DotPlots of relevant marker genes used to annotate cell populations in human (C) and mouse (D). Dot size represents the percentage of cells expressing the marker gene within a population, and color scale indicates normalized expression level. (E and F) UMAP colored by level of expression of representative supporting markers (*FOXL2/Foxl2*, *BMP2/Bmp2*, *CYP19A1, LHX2, Runx1, Lgr5)* and stromal markers (*NR2F2, Wnt5a, Arx, Pdgfra*) in human (E) and mouse (F) datasets. (G and H) Barplots representing the weights of the top 50 genes of supporting or SP identity models in human (G) and mouse (H). (I and J) Stage-wise UMAP representations colored by cell type with cPAGA connectivity between cell types in human (I) and mouse (J). Cell type abbreviations: CE, coelomic epithelium; SE, surface epithelium; SLC, supporting-like cells; pre-Sup, pre-supporting cells; GM, gonadal mesenchyme; pre-Gran, pre-granulosa cells; Gran, granulosa cells; SP, stromal progenitors; Stero, steroidogenic cells; SM, smooth muscle cells; PV, perivascular cells; Luteal, luteal cells; MT, mesonephric tubules; MM, mesonephric mesenchyme; Germ, germ cells; AS, Adreno Sympathic ; End, endothelial cells; Eryt, erythrocytes; Immune, immune cells.

**Supplementary Figure S2.**
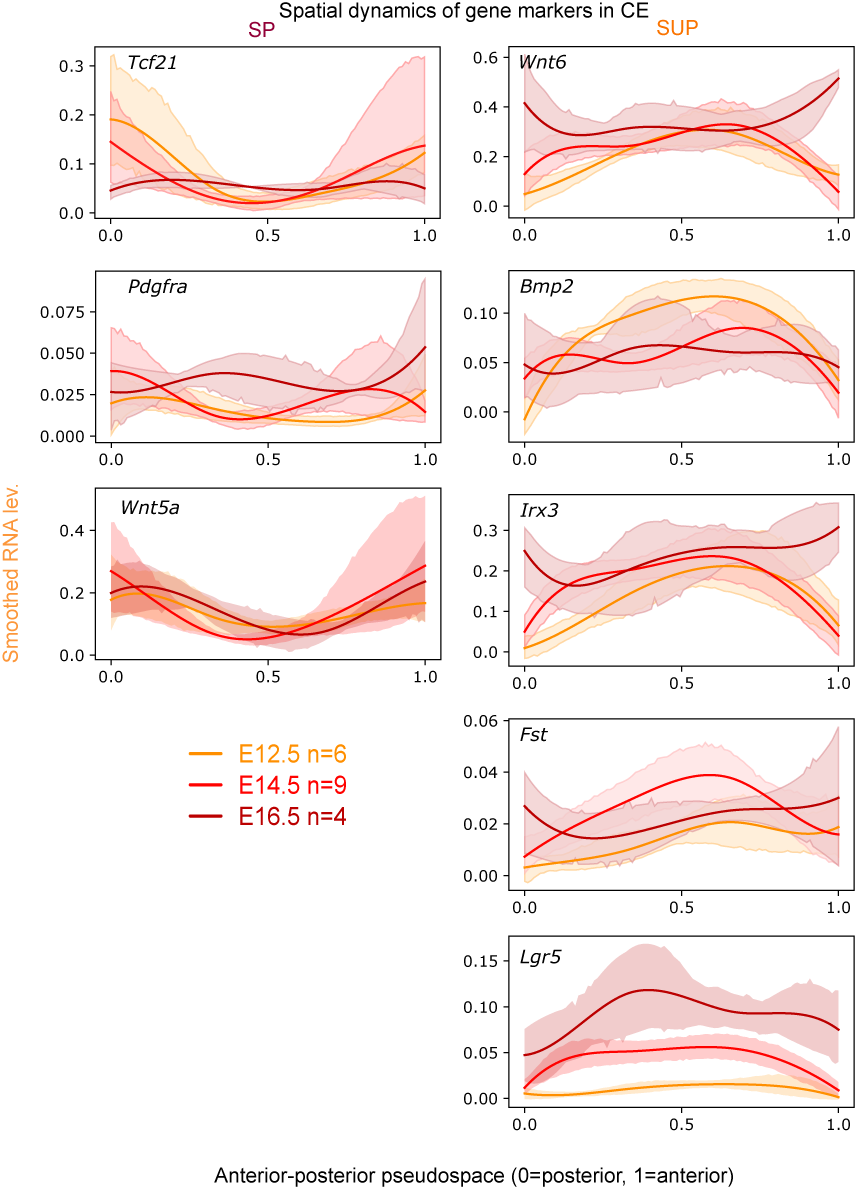
related to Figure 2. Lineplots of smoothed expression level of stromal markers (*Tcf21, Pdgfra, Arx* and *Wnt5a*) and supporting markers (*Wnt6*, *Bmp2*, *Irx3*, *Lgr5* and *Fst*) along CE pseudospace at E12.5, E14.5 and E16.5 (Mean with 95% confidence intervals). In the pseudospace representation, the posterior pole of the ovary is set at 0 and the anterior pole at 1.

**Supplementary Figure S3.**
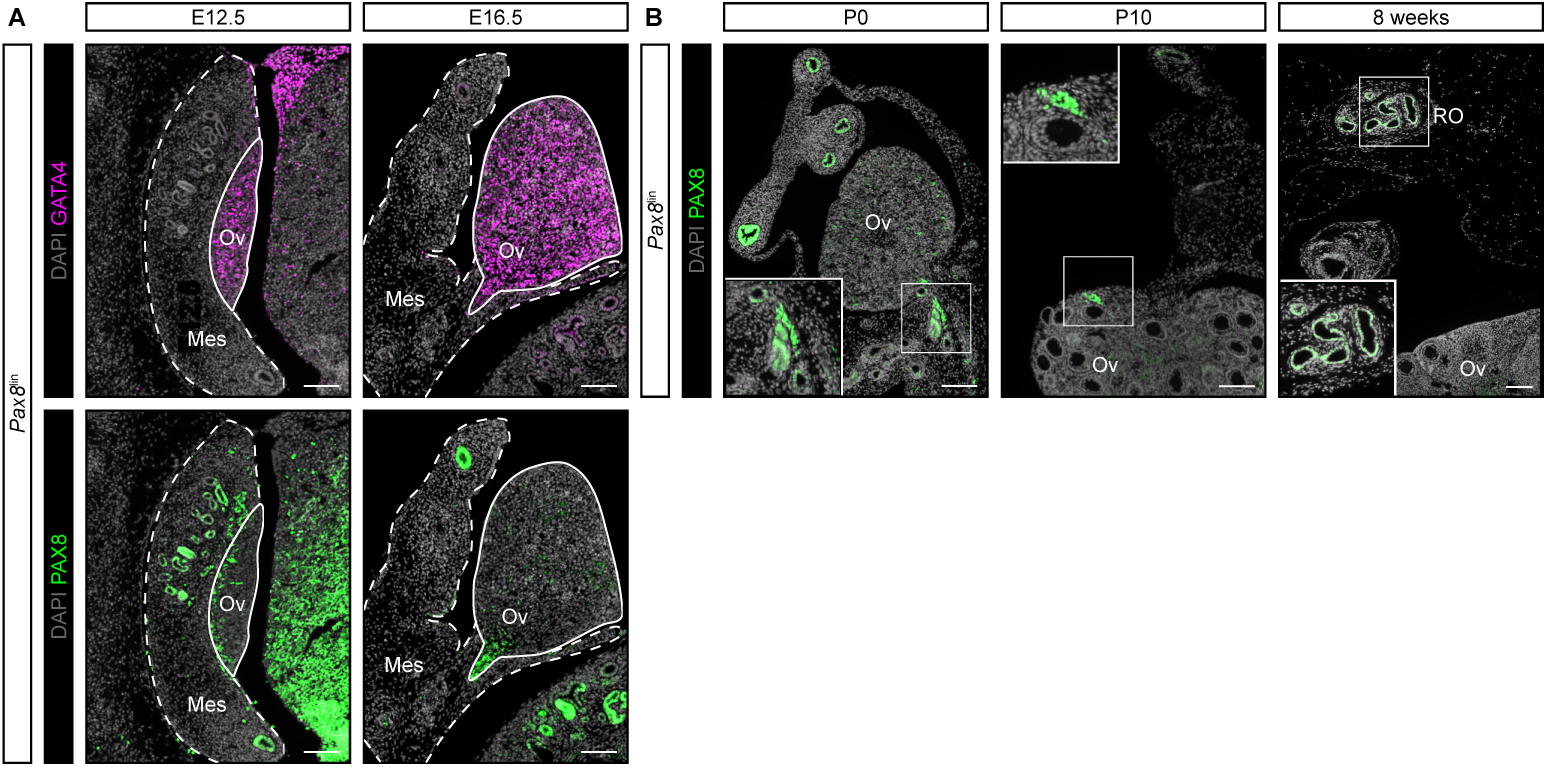
related to Figure 3. (A) Co-IF for GATA4 and PAX8 in E12.5 and E16.5 fetal ovaries. White line: ovarian domain, dashed line: mesonephric domain. (B) IF for PAX8 in P0, P10 and adult *Pax8*^lin^ ovaries showing PAX8 expression restricted to the rete ovarii.

## Notes

### Competing Interest Statement

The authors have declared no competing interest.

### Summary of Updates

This revised version is more concise and clarifies the key messages. We have added human scRNA-seq data demonstrating that the mechanisms observed in mice are conserved in humans, albeit with different temporal dynamics. We also include new ex vivo culture experiments highlighting the influence of the mesonephros on coelomic epithelial cell fate during ovarian development. Accordingly, the Results section and figures have been expanded, and additional collaborators have been included as authors.

